# Molecular and morphological analyses clarify species delimitation and reveal a new *Betula* species in section *Costatae*

**DOI:** 10.1101/2020.10.29.361519

**Authors:** Luwei Wang, Junyi Ding, James S. Borrell, Hugh A. McAllister, Feifei Wang, Lu Liu, Nian Wang

## Abstract

**Background and Aims:** Delineating closely related and morphologically similar species with overlapping ranges can be difficult. Here, we use section *Costatae* (genus *Betula*) as a model to resolve species and subspecies boundaries in four morphologically similar trees: *Betula ashburneri, Betula costata, Betula ermanii* and *Betula utilis* (including ssp. *utilis,* and diploid and tetraploid races of ssp. *albosinensis*).

**Methods:** We genotyped 298 individuals (20-80 per species) from 38 populations at 15 microsatellite markers and a subset of 34 individuals from 21 populations using restriction-site associated DNA sequencing (RAD-seq). Morphometric analysis was conducted to characterise leaf variation for a subset of 89 individuals.

**Key Results:** Molecular analyses and leaf morphology found little differentiation between *B. ashburneri,* diploid *B. utilis* ssp. *albosinensis* and some samples of *B. utilis* ssp. *utilis* suggesting that these should be treated as a single species. By contrast, tetraploid *Betula utilis* ssp. *albosinensis* was divided into two groups with group I genetically similar to *B. utilis* ssp. *utilis* based on SNPs and group II, a very distinct cluster, which we propose as a new species, namely, *Betula buggsii*. Phylogenomic analysis based on 2,285,620 SNPs show a well-supported monophyletic clade of *B. buggsii,* forming a sister with a well-supported clade of *B. ashburneri,* diploid *B. albosinensis* and some samples of *B. utilis* ssp. *utilis*. Morphologically, *Betula buggsii* is characterised by elongated lenticels and a distinct pattern of bark peeling. *Betula buggsii* is geographically restricted to the Qinling-Daba Mountains.

**Conclusions:** Our study reveals six genetically distinguishable species: *B. ashburneri, B. buggsii, B. costata, B. utilis* ssp. *utilis*, *B. utilis* ssp. *albosinensis* and *B. ermanii*. Our research demonstrates an integrative approach in delimitating species using morphological and genetic samples from their nearly entire distributions. Analyses based on subsets of species’ distributions may lead to erroneous species or subspecies delineation.

## INTRODUCTION

Species delineation based on morphology may be confounded by intra-specific variation among populations and limited differentiation between closely-related species (Whittall et al., 2004; Leliaert et al., 2009; Wang et al., 2014b; Lissambou et al., 2019). Where species co-occur, this may be further exacerbated by introgression and hybridisation (Bacon et al., 2012; Andújar et al., 2014), or may be morphologically impossible due to cryptic speciation (Bickford et al., 2007; Fišer et al., 2018). Despite advances in phylogenetic methods, this has meant that many species rich genera have remained unresolved, hindering our understanding of species ecology and evolution as well as limiting our ability to deliver effective conservation management.

*Betula* L. (Betulaceae) is such a genus with many taxonomic issues. The genus consists of approximately 65 species and subspecies (Ashburner & McAllister, 2016) with some spanning a very broad latitudinal and longitudinal range, such as *B. platyphylla,* ranging from Europe to eastern Asia and from the Himalayas to Siberia. Species such as *B. michauxii* and *B. nana,* are morphologically convergent but comparatively distantly-related (Wang et al., 2016; Wang et al., 2020), having evolved independently in North America and Scotland respectively. Analysis of *Betula* taxonomy is complicated by these broad ranges, frequent inter-specific hybridisation (Anamthawat-Jónsson & Tómasson, 1999; Wang et al., 2014a; Zohren et al., 2016; Tsuda et al., 2017), polyploidy and considerable morphological variation (Wang et al., 2014b; Ashburner & McAllister, 2016).

Where ranges overlap, introgression appears frequent between species of the same ploidy level (Nagamitsu et al., 2006; Ashburner & McAllister, 2016) and even differing ploidy levels (Anamthawat-Jónsson & Thórsson, 2003; Wang et al., 2014a; Zohren et al., 2016; Tsuda et al., 2017). *Betula* species from different subgenera appear able to hybridise readily, such as hybridisation between *B. alleghanensis* and *B. papyriflera* (Thomson et al., 2015). Polyploidy is also common within *Betula,* accounting for nearly 60% of the described taxa, ranging from diploid to dodecaploidy, with cytotypes observed for some species, such as *B. chinensis* (6x and 8x) (Ashburner & McAllister, 2016).

In this study, we use section *Costatae* as a model in which to demonstrate combined morphological and genetic methods to resolve these taxonomic issues (Table 1). Section *Costatae* includes the diploids *B. ashburneri* and *B. costata,* with *B. ashburneri* discovered from south-east Tibet and reported to have distributions in north-west Yunnan and western Sichuan (McAllister & Rushforth, 2011) and with *B. costata* distributed in northern and northeastern China, Japan and Russian Far East (Ashburner & McAllister, 2016). Section *Costatae* also includes two tetraploids: *B. utilis* (subdivided into ssp. *utilis* and ssp. *albosinensis*) occurring from the Himalayas to north China without clear geographical and morphological intra-specific boundaries and *B. ermanii* from northeastern China, Japan and Russian Far East. Several varieties of these tetraploid species have also been named based on a limited number of herbarium specimens, such as *B. utilis* var. *prattii, B. albosinensis* var. *septentrionalis* and *B. ermanii* var. *lanata* (Ashburner & McAllister, 2016), though their taxonomic validity is unclear.

**Table 1.** Detailed information on taxa of section *Costatae* used in the present study.

| Species | Variety | Ploidy | Distribution | Reference |
| --- | --- | --- | --- | --- |
| *B. utilis* | *ssp. utilis* | tetraploid | Nepal, eastwards through SE Tibet to Yunnan and west Sichuan where it merges with *ssp. albosinensis* | Ashburner and McAllister, 2016 |
|  | *ssp. utilis* | tetraploid | Gansu, Hebei, Ningxia, Qinghai, Shaanxi, west Sichuan, E and S Xizang, NW Yunnan | Li and Alexei, 1999, Flora of China |
|  | *var. pratii* | tetraploid | Kangding, western Sichuan | Ashburner and McAllister, 2016 |
|  | *ssp. albosinensis* | tetraploid | North Sichuan, Hubei, south Gansu, south Ningxia, south Shaanxi, Shanxi, Henan and Hebei | Ashburner and McAllister, 2016 |
|  | *ssp. albosinensis* | diploid | south Shaanxi | Hu et al., 2019 |
| *B. albosinensis* | *ssp. septentrionalis* | tetraploid | Western Sichuan | Ashburner and McAllister, 2016 |
| *B. ermanii* | *ssp. ermanii* | tetraploid | Northeast China, Japan, Korea and the Russian Far East | Ashburner and McAllister, 2016 |
|  | *var. lanata* | tetraploid | Russia: from the eastern shores of Lake Baikal, eastward to the Pacific coast except Korea and north China | Ashburner and McAllister, 2016 |
| *B. costata* | NA | diploid | Northeast China | Ashburner and McAllister, 2016 |
| *B. ashburneri* | NA | diploid | Southeast Tibet, Northwest Yunnan, Southwest Sichuan and possibly Shaanxi | McAllister, 2011; Ashburner and McAllister, 2016 |

Confusingly, *B. utilis* ssp. *utilis* was described to have distributions in Gansu, Ningxia, Qinghai and Shaanxi according to Flora of China (Li & Skvortsov, 1999), where *B. utilis* ssp. *utilis* was not recorded in Ashburner and McAllister’s monograph (Ashburner & McAllister, 2016) (Table 1). Recently, a ‘diploid’ *B. albosinensis* has been discovered from the Qinling Mountains in central China (Hu et al., 2019). A phylogenetic tree based on the internal transcribed spacer (ITS) region indicated a close relationship between the ‘diploid’ *B. albosinensis, B. ashburneri* and *B. costata* (Hu et al., 2019).

It remains unknown if the ‘diploid’ *B. albosinensis, B. ashburneri* and *B. costata* represent distinct genetic entities. Moreover, it remains unknown if the tetraploids *B. utilis* ssp. *albosinensis, B. ermanii* and *B. utilis* ssp. *utilis* described in Flora of China and in Ashburner and McAllister’s monograph, respectively, represent distinct genetic entities (Li & Skvortsov, 1999; Ashburner & McAllister, 2016). For ease of reference, we abbreviate the ‘diploid’ *B. albosinensis* and *B. utilis* ssp. *utilis* described in Ashburner and McAllister’s monograph and in Flora of China as *B. albosinenesis* [DA], *B. utilis* [AM] and *B. utilis* [FC], respectively.

To resolve taxonomic issues within section *Costatae,* we carried out morphological analysis, microsatellite genotyping and restriction-site associated DNA sequencing (RAD-seq). Our specific aims are to (1) identify the number of distinct genetic groups within section *Costatae,* with particular attention to (2) resolving the phylogenetic position of *B. utilis* [FC] and *B. albosinensis* [DA]; finally, (3) we integrate genetic data with morphology and geographic distributions to present a revised treatment of species boundaries within section *Costatae*. We consider the applicability of this approach to other taxonomically complex genera.

## Materials and methods

### Sampling

Samples putatively identified (based on morphology) as *B. utilis* [AM], *B. utilis* ssp. *albosinensis, B. ermanii, B. albosinensis* [DA], *B. utilis* [FC] and *B. costata* were collected from between three and twelve populations each (Fig. 1). All species were collected from naturally occurring woodland, meaning that they were not artificially planted. Leaf samples were collected between May and September of 2018 and 2019, with each sample separated by ~20m. A herbarium specimen was created for each sample except for a subset of samples where branches were difficult to obtain. For these samples, cambium tissue was collected. A GPS system (UniStrong) was used to record the coordinate points of each population. Detailed species and population sampling information is provided in Supplementary Data Table S1.

**Figure 1.**
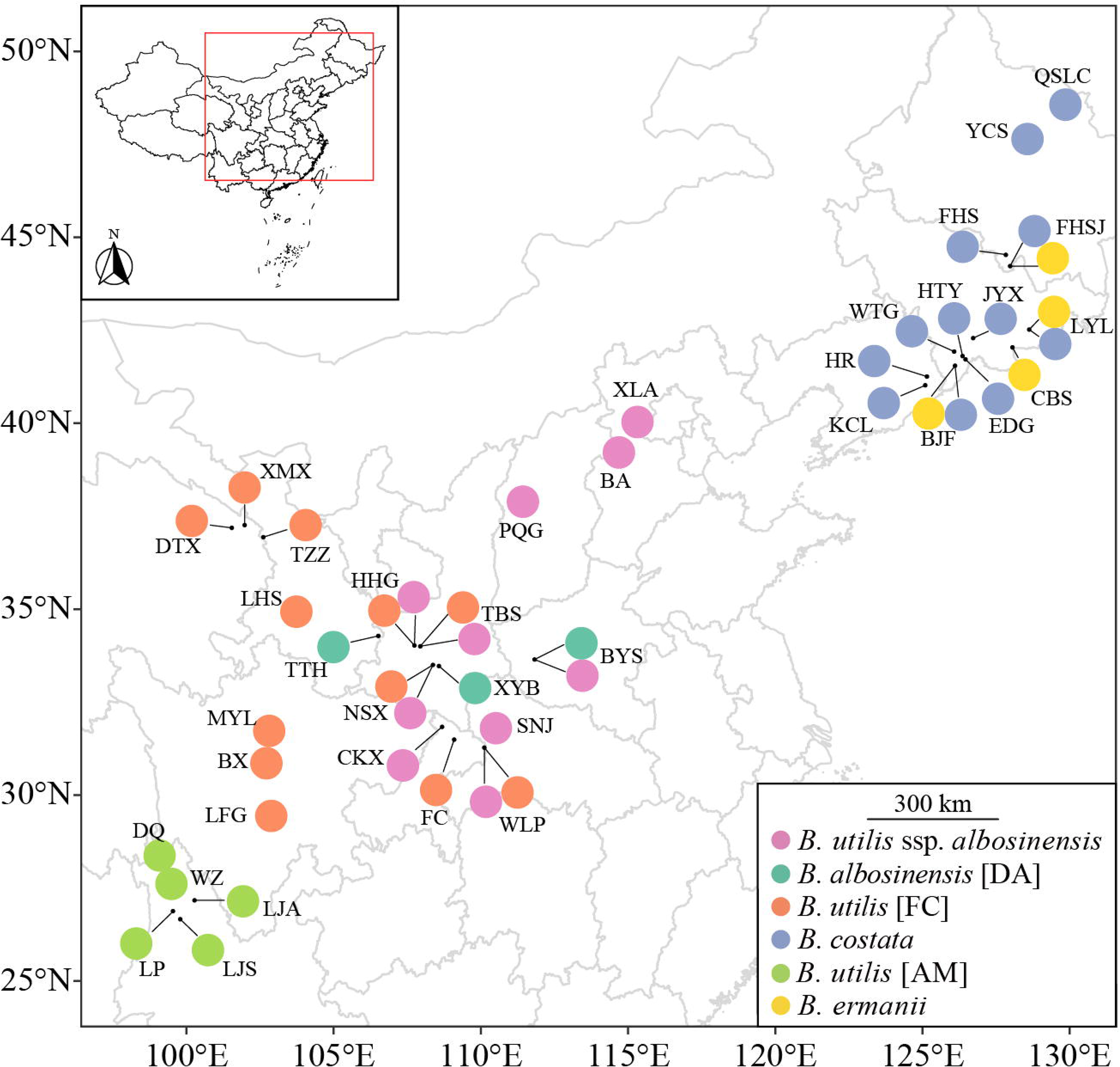
The distribution of samples used in the present study.

### Species identification

*Betula utilis* [AM] is distributed through SE Tibet to Yunnan and Sichuan and *B. utilis* ssp. *albosinensis* occurs in North Sichuan, Hubei, Shaanxi, Shanxi, Henan and Hebei (Ashburner and McAllister, 2016). These two species co-occur in Sichuan province. Due to a morphological continuum between *B. utilis* [AM] and *B. utilis* ssp. *albosinensis,* we assigned our populations based on geographic origins, with populations from northwestern Yunnan designated as *B. utilis* [AM] and populations from south Shaanxi, Hubei, Shanxi, Henan and Hebei designated as *B. utilis* ssp. *albosinensis. Betula utilis* [FC] occupies a higher altitude than *B. utilis* ssp. *albosinensis* and can be distinguished from the latter by its leathery dark green leaves (Ashburner & McAllister, 2016; Li & Skvortsov, 1999). *Betula costata* and *B. ermanii,* having distributions in northeastern China, can be distinguished from leaf morphology with the former having lanceolate leaves and the latter triangular-ovate leaves (Li & Skvortsov, 1999; Ashburner & McAllister, 2016).

### Morphometric analyses

For analyses of leaf shape among these species, we selected 6-27 individuals per taxa and sampled 283 leaves. Leaves were scanned individually using a Hewlett-Packard printer (LaserJet Pro MFP M128fn) with a resolution of 600 dpi. Thirteen landmarks were selected from each scanned leaf according to the protocols of (Liu et al., 2018; Hu et al., 2019). The 13 landmarks were converted to a configuration of 26 cartesian coordinates using ImageJ (Abràmoff et al., 2004). A Generalized Procrustes Analysis (GPA) was performed using the procGPA function in the R package “shapes” (Dryden, 2019). Eigenleaves were visualized using the “shapepca” function and principal component scores, percentage variance and Procrustes-adjusted coordinates were obtained from procGPA object values.

### DNA extraction and microsatellite genotyping

High quality DNA was extracted from cambial tissues following a modified 2x CTAB (cetyltrimethylammonium bromide) protocol (Wang et al., 2013). Extracted DNA was assessed with 1.0% agarose gels. Fifteen microsatellite loci developed for *B. platyphylla* var. *japonica* (Wu et al., 2002), *B. pendula* (Kulju et al., 2004), *B. pubescens* ssp. *tortuosa* (Truong et al., 2005) and *B. maximowicziana* (Tsuda et al., 2009) were used to genotype our samples (Supplementary Data, Table S2), with the 5’ terminus of the forward primers labeled with FAM, HEX or TAM fluorescent probes. These microsatellite loci have a good cross compatibility in multiple *Betula* species. Each microsatellite locus was amplified individually and was artificially combined into four multiplexes. The PCR protocol followed Hu et al. (2019). Microsatellite alleles were scored using GENEMARKER 2.4.0 (Softgenetics) and checked manually. Individuals with more than three missing loci were excluded for further analyses, resulting in 298 individuals in the final dataset.

### RAD-seq

A subset of 34 DNA samples were selected for RAD-seq using an Illumina HiSeq 2500 and 150-bp pair-end sequencing with the restriction enzyme *PstI* (Personalbio company, Shanghai, China). These were combined with eight additional samples of section *Costatae* previously sequenced, using the same restriction enzyme in Wang et al. (2020). These samples represented six *B. costata,* six *B. utilis* [AM], six *B. ermanii,* twelve *B. utilis* ssp. *albosinensis,* seven *B. utilis* [FC] and one of each of *B. albosinensis* [DA], *B. ashburneri, B. ermanii* var. *lanata, B. albosinensis* var. *septentrionalis*, and *B. utilis* var. *prattii* (Supplementary Data, Table S3). The raw data were trimmed using Trimmomatic (Bolger et al., 2014) in paired-end mode. Reads with a quality of below 20 within the sliding-window of 5 bp and unpaired reads were discarded. We performed LEADING and TRAILING to remove bases with a quality below 20. Then we performed a SLIDINGWINDOW step to discard reads shorter than 40 bp. Filtered reads of each sample were aligned to the whole genome sequence of *B. pendula* (Salojärvi et al., 2017) using BWA-MEM v.0.7.17-r1188 algorithm in BWA (v0.7.17) with default parameters (Li & Durbin, 2009). Non-specific mapped reads were discarded. All subsequent analyses were performed using SAMtools v1.8 (Li et al., 2009) and GATK V4.1.4 (McKenna et al., 2010; DePristo et al., 2011). These include conversion of alignments into indexed binary alignment map (BAM) files, marking duplicates, calling genotypes and filtering SNPs (McKenna et al., 2010; DePristo et al., 2011). SNPs within a 50 kb window with r^2^ > 0.5 and a minimum allele frequency (MAF) < 0.01 were removed to reduce linkage disequilibrium using BCFtools v1.10.2 (Li, 2011). Prior to population structure analysis, we retained only sites with no missing data, resulting in 82,137 SNPs for downstream analyses.

### Analyses of microsatellite data and SNPs

A principal coordinate analysis (PCoA) was performed on microsatellite data of *B*.

*utilis* [AM], *B. utilis* [FC], *B. utilis* ssp. *albosinensis, B. albosinensis* [DA], *B. costata* and *B. ermanii* using POLYSAT (Clark & Jasieniuk, 2011) implemented in R 4.0.2 (R Core Team, 2020), based on Bruvo’s genetic distances (Bruvo et al., 2004). For nucleotide SNPs, a principal component analysis (PCA) was carried out using the ‘adegenet’ R package 2.1.1 (Jombart, 2008).

Microsatellite data were analyzed in STRUCTURE (Pritchard et al., 2000) to identify the most likely number of genetic clusters (K) with a ploidy of four. Ten replicates were performed with 1,000,000 iterations and a burn-in of 100,000 for each run at each value of K from 1 to 8. We used the admixture model, with an assumption of correlated allele frequencies among populations. Individuals were assigned to clusters based on the highest membership coefficient averaged over the ten independent runs. The number of genetic clusters was estimated using the “Evanno test” (Evanno et al., 2005) implemented in Structure Harvester (Earl & vonHoldt, 2012). Replicate runs were grouped based on a symmetrical similarity coefficient of >0.9 using the Greedy algorithm in CLUMPP (Jakobsson & Rosenberg, 2007) and visualized in DISTRUCT 1.1 (Rosenberg, 2004).

The filtered SNPs were analyzed in ADMIXTURE v1.3.0, a model-based approach to assessing population structure in a Maximum Likelihood framework (Alexander & Lange, 2011). We ran ADMIXTURE for K = 1-10 with 20 replicates for each K value and performed cross-validation error estimation in order to assess the most suitable value of K (Alexander & Lange, 2011). Replicate runs were aligned and visualised in pong v1.4.9 with the greedy algorithm (Behr et al., 2016).

### ITS and SNP based phylogenetic analyses

To provide an additional line of evidence for the phylogenetic position of *B. utilis* [FC], *B. albosinensis* [DA], and *B. costata,* we generated ITS sequence and SNP based phylogenies.

First, we amplified the nuclear ribosomal internal transcribed spacer (nrITS) region (ITS1, 5.8S and ITS2) using primers ITS4 (White et al., 1990) and ITSLeu (Baum et al., 1998), with seven, ten, five and four individuals of *B. utilis* ssp. *albosinensis* group II collected from the NSX, CKX, WLP and SNJ, respectively. The reaction mix and the PCR protocol followed that of Hu et al. (2019). PCR products were purified and sequenced at Tsingke Company (Qingdao, China). Sixty-four additional ITS sequences from Betulaceae (Wang et al., 2016) were included to infer the phylogenetic position of *B. utilis* ssp. *albosinensis* group II. In total, 90 sequences were aligned using BioEdit v7.0.9.0 (Hall, 1999) with default parameters.

Second, we collated RAD-seq data of 20 *Betula* taxa representing genus wide diploid species. The identity of the 20 sequenced *Betula* taxa was initially inferred via ITS sequences and genome size estimates (Wang et al., 2016). In addition, we included RAD-seq data of 17 samples generated in the present study. *Alnus inokumae* was selected as the outgroup (Supplementary Data, Table S3). SNPs of a total of 38 taxa were concatenated into a supermatrix for phylogenetic analysis. SNPs with a missing data > 50% were excluded, resulting in 2,285,620 SNPs.

For both the ITS alignment and the matrix of SNPs, we conducted a rapid bootstrap analysis under a GTR+GAMMA nucleotide substitution model, with 100 bootstraps and 10 searches using the maximum-likelihood method (ML) in RAxML v. 8.1.16 (Stamatakis, 2006). The phylogenetic trees were visualised in FigTree v.1.3.1.

## Results

### Morphometric analyses

Landmarks were first aligned using a GPA and then a principal component analysis (PCA) was conducted to visualise the major sources of shape variance of leaves from *B. albosinensis* [DA], *B. utilis* ssp. *albosinensis, B. utilis* [AM], *B. utilis* [FC], *B. costata* and *B. ermanii*. PC1 and PC2 produce largely overlapping clusters among *B. albosinensis* [DA], *B. utilis* ssp. *albosinensis, B. utilis* [AM] and *B. utilis* [FC], but *B. costata* and *B. ermanii* overlapped to a much lesser extent (Fig. 2a). The shape variance, represented by PC1 and PC2, is mainly influenced by leaf width and marginally influenced by leaf length (Fig. 2b).

**Figure 2.**
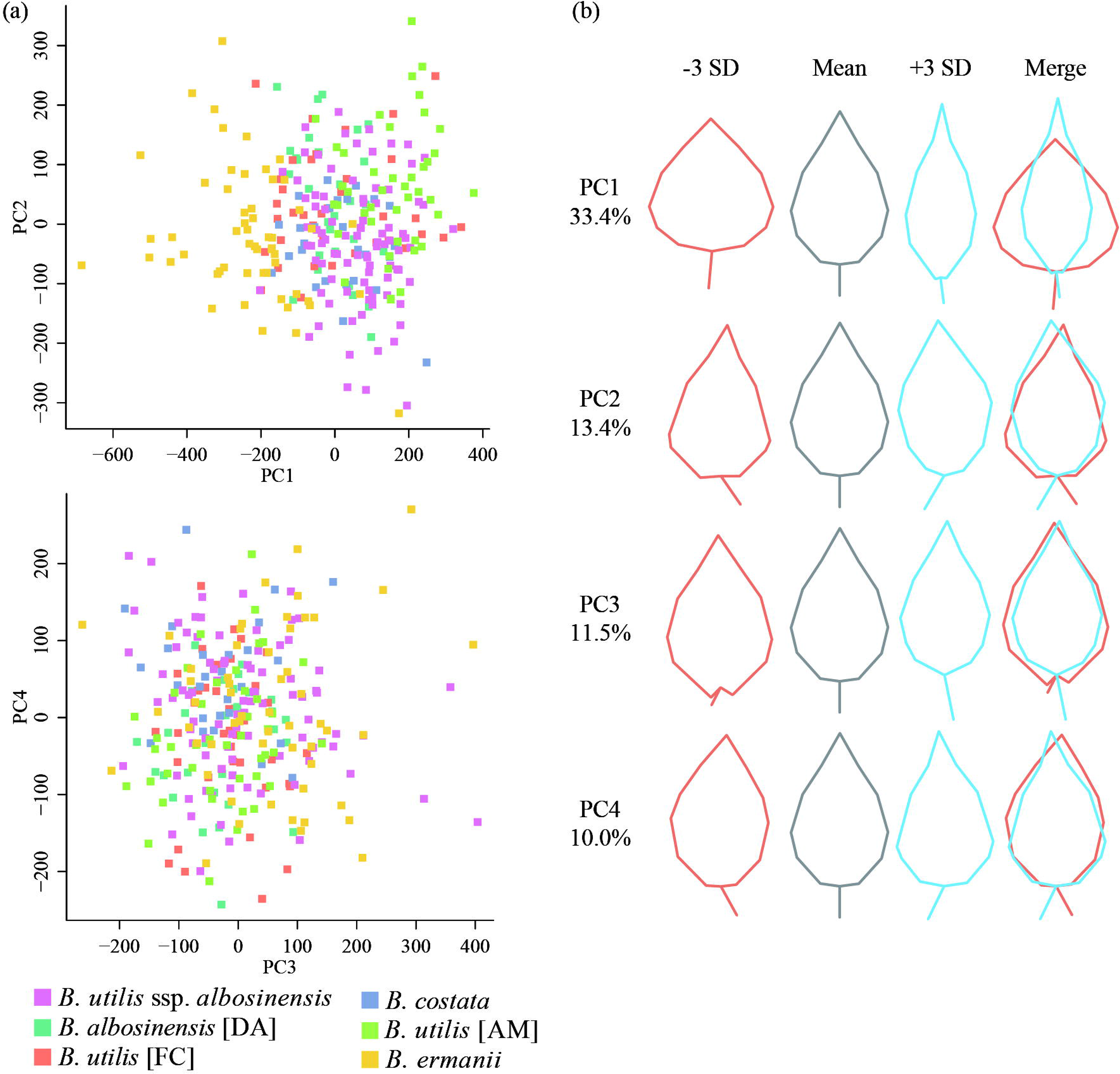
(a) Principal component analysis (PCoA) of leaves of section *Costatae* species. (b) ‘Eigenleaves’ showing leaf morphs represented by principal components (PCs) at ± 3SD and shape variance explained by each PC. Each dot represents a leaf.

### PCO and PCA analyses

PCO analysis based on microsatellite markers revealed five clusters, with the first three axes accounting for 43.8% of the total variation (Fig. 3a). *Betula utilis* ssp. *albosinensis* forms two groups: group I overlaps substantially with *B. utilis* [AM] and *B. ermanii* whereas group II separates from all the other species on coordinate 1 (Fig. 3a). *Betula utilis* [FC] and *B. albosinensis* [DA] overlap substantially whereas *B. costata* separates from the remaining species on coordinates 2 and 3 (Supplementary Data, Fig. S1a). *Betula ermanii* separates from *B. utilis* [AM] on coordinate 3 with *B. utilis* ssp. *albosinensis* group I intermediate (Supplementary Data, Fig. S1a).

**Figure 3.**
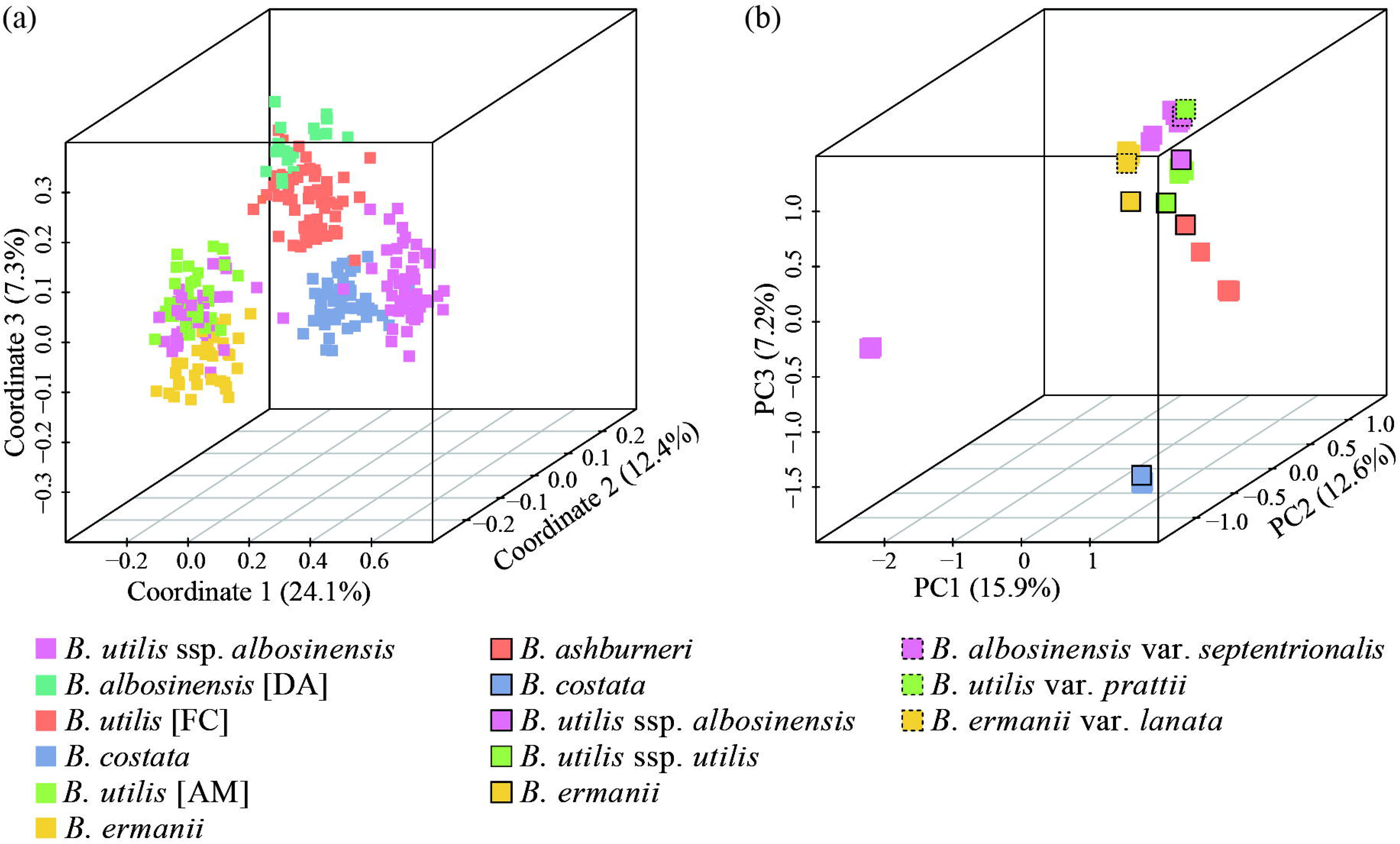
Principal coordinate analysis (PCO) of section *Costatae* species at 15 microsatellite markers (a) and principal component analysis (PCoA) of section *Costatae* species at 82,137 SNPs (b).

For the sequenced individuals, between 12,234,848 and 28,155,092 reads were retained for each individual (mean 18,862,242) after trimming and filtering (Supplementary Data, Table S3). The number of variable sites of the sequenced individuals ranges from 5,520,333 to 9,735,507. A principal component analysis (PCA) based on genotype calls for 82,137 SNPs shows that both *B. utilis* ssp. *albosinensis* group II and *B. costata* separate from the remaining species and from each other (Fig. 3b). Two individuals of *B. utilis* ssp. *albosinensis* group I form a cluster and three individuals of *B. utilis* ssp. *albosinensis* group I form a cluster with the previously sequenced *B. utilis* var. *prattii* and *B. albosinensis* var. *septentrionalis* (Fig. 3b). *Betula utilis* [AM] forms a cluster with the previously sequenced *B. utilis* ssp. *albosinensis* whereas *B. ermanii* and *B. ermanii* var. *lanata* form a cluster from PC1 and PC2 (Supplementary Data, Fig. S1b). *Betula albosinensis* [DA] forms a cluster with one accession of *B. utilis* [FC] whereas the remaining accessions of *B. utilis* [FC] form another cluster. The two individuals of *B. utilis* ssp. *albosinensis* group I position between *B. ermanii* and three individuals of *B. utilis* ssp. *albosinensis* group I. *Betula ashburneri* forms a continuum with *B. utilis* [AM] and *B. utilis* [FC] on PC1 and PC3 (Supplementary Data, Fig. S1b).

### STRUCTURE and ADMIXTURE analyses

STRUCTURE analyses based on microsatellite markers identified five clusters: (1) *B. utilis* ssp. *albosinensis* group I, (2) *B. albosinensis* [DA] and *B. utilis* [FC], (3) *B. costata,* (4) *B. utilis* [AM], (5) *B. ermanii* and (6) *B. utilis* ssp. *albosinensis* group II (Supplementary Data Figs. S2-3). *B. utilis* ssp. *albosinensis* group I is genetically similar to *B. ermanii* at all K values (Fig. 4a). *Betula utilis* ssp. *albosinensis* group II includes populations SNJ, WLP, NSX and CKX and separates with the remaining species (Supplementary Data, Fig. S3). Similarly, *B. albosinensis* [DA] and *B. utilis* [FC] are genetically similar at all supported K values (Supplementary Data, Fig. S3). Admixture analysis based on the same set of SNPs showed that cross-validation error is smallest at K = 5, but only with four out of twenty replicates having an average pairwise similarity of 0.98 (Supplementary Data, Fig. S4). At the value of K = 6, fourteen out of twenty replicates have an average pairwise similarity of 0.98. However, the cross-validation error is slightly larger than that when K = 5 (Supplementary Data, Fig. S4). At the value of K = 5, *B. utilis* ssp. *albosinensis* group I genetically resembles *B. utilis* [AM] with exception of samples XLA01 and XLA32, which are more genetically similar to *B. ermanii* (Fig. 4b). At the value of K = 6, *B. utilis* ssp. *albosinensis* group I separates from *B. utilis* [AM] and XLA01 and XLA32 exhibit genetic admixture from *B. ermanii* (Fig. 4b). *Betula utilis* ssp. *albosinensis* group II separates from the remaining species at the value of K = 3 and onwards (Supplementary Data, Fig. S5). Interestingly, this identified that *B. albosinensis* var. *septentrionalis* and *B. utilis* var. *prattii* are genetically similar to *B. utilis* ssp. *albosinensis* group I whereas the *B. utilis* ssp. *albosinensis* and *B. utilis* ssp. *utilis* are genetically similar to *B. utilis* [AM] (Fig. 4b).

**Figure 4.**
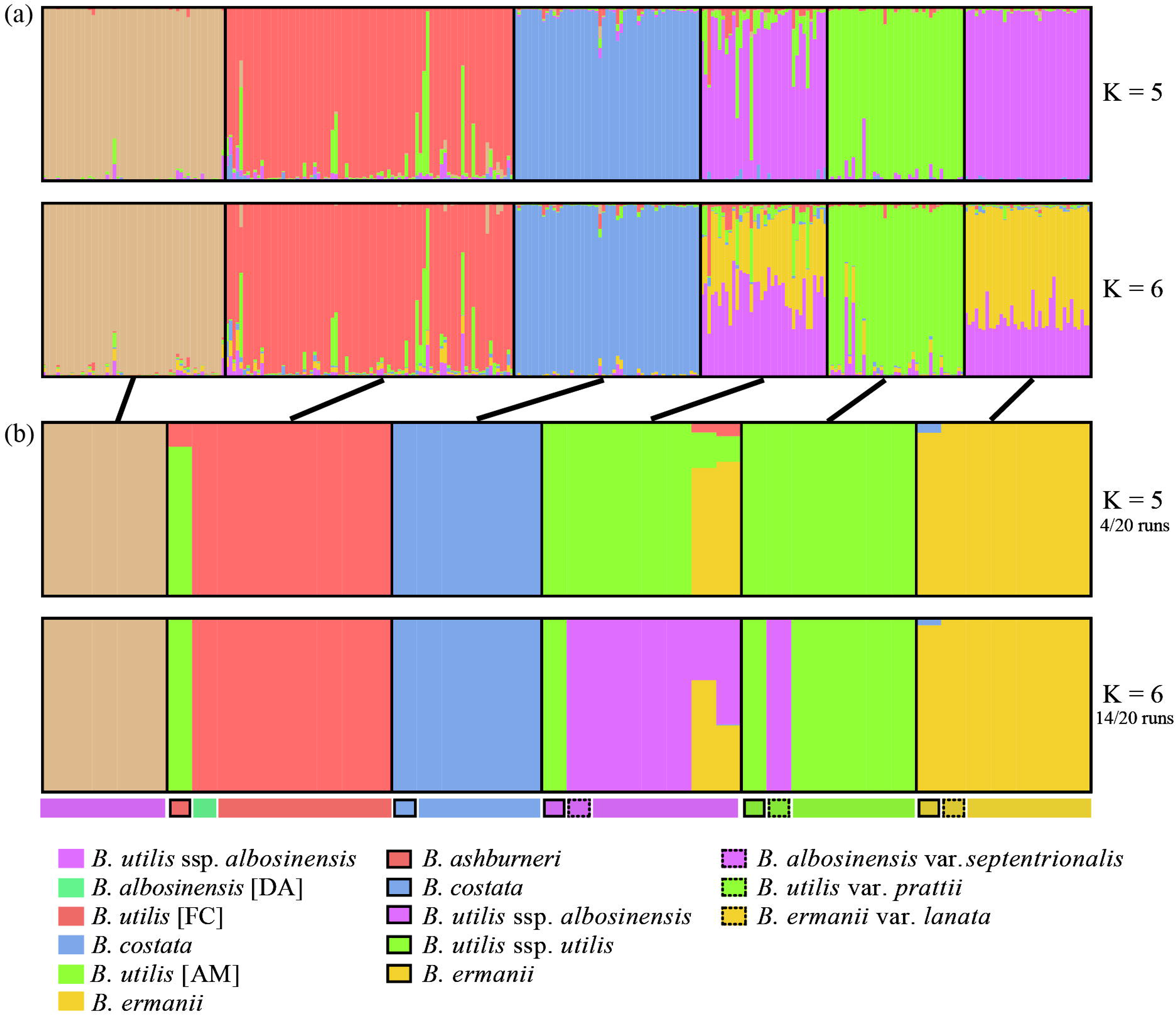
STRUCTURE results of section *Costatae* at K values 5 and 6 based on 15 microsatellite markers (a) and admixture analysis of section *Costatae* at K values 5 and 6 at the 82,137 SNPs (b).

### Phylogenetic analyses

#### Identification of species novo *Betula buggsii*

Analyses of microsatellite data and SNPs indicate that *B. utilis* ssp. *albosinensis* group II is genetically distinct from other species of section *Costatae* and therefore represents a putative new species, namely *Betula buggsii*. Morphologically, despite general similarity to *B. utilis* ssp. *albosinensis,* we found *B. buggsii* is characterised by very elongated lenticels with bark peeling along lenticels into strips.

The phylogenetic tree based on ITS showed that *B. buggsii* samples formed a monophyletic cluster, within a clade with species of section *Acuminatae, B. bomiensis* and *B. nigra*. However, this clade received little support (Supplementary Data, Fig. S6). The phylogenetic tree based on a matrix of 2,285,620 SNPs showed that the five individuals of *B. buggsii* from populations CKX, SNJ and WLP formed a monophyletic clade with 100% support, which was basal to a clade of *B. costata, B. ashburneri, B. albosinensis* [DA] and *B. utilis* [FC] (Fig. 5). The five individuals of *B. costata* formed a monophyletic clade whereas individuals of *B. utilis* [FC], *B. albosinensis* [DA] and *B. ashburneri* intermixed and together formed a monophyletic clade with 100% support (Fig. 5).

**Figure 5.**
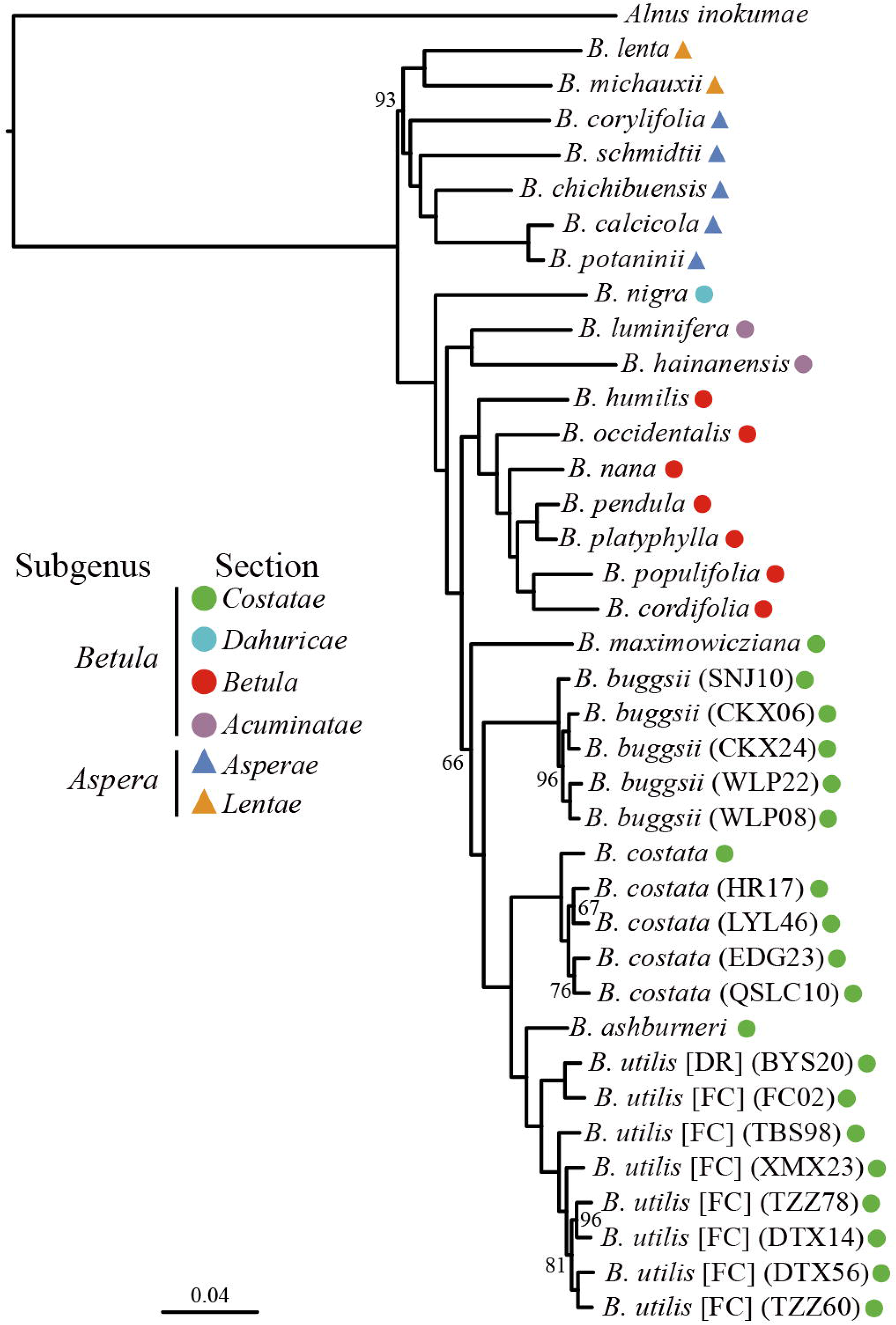
Species tree from the maximum likelihood analysis of the diploid *Betula* species using the supermatrix approach based on data from 2,285,620 SNPs. Bootstrap support values of 100 were not shown. Numbers on the branches are bootstrap support values between 60 and 100. The scale bar below indicates the mean number of nucleotide substitutions per site. Species were classified according to Wang et al. (2020).

#### Taxonomic treatment

##### *Betula buggsii* N. Wang, sp. nov

###### Diagnosis

*Betula buggsii* is very similar with *B. utilis* ssp. *albosinensis* in leaf morphology but have very elongated lenticels on the bark of adult and elderly trees. Barks of *B. buggsii* peel along the elongated lenticels into strips. Adult or old *B. utilis* ssp. *albosinensis* trees exfoliate in large sheets (Fig. 6a). Seedlings of *B. buggsii* and *B. utilis* ssp. *albosinensis* (DBH < 5 cm) show no obvious difference in bark color and patterns of bark peeling.

**Figure 6.**
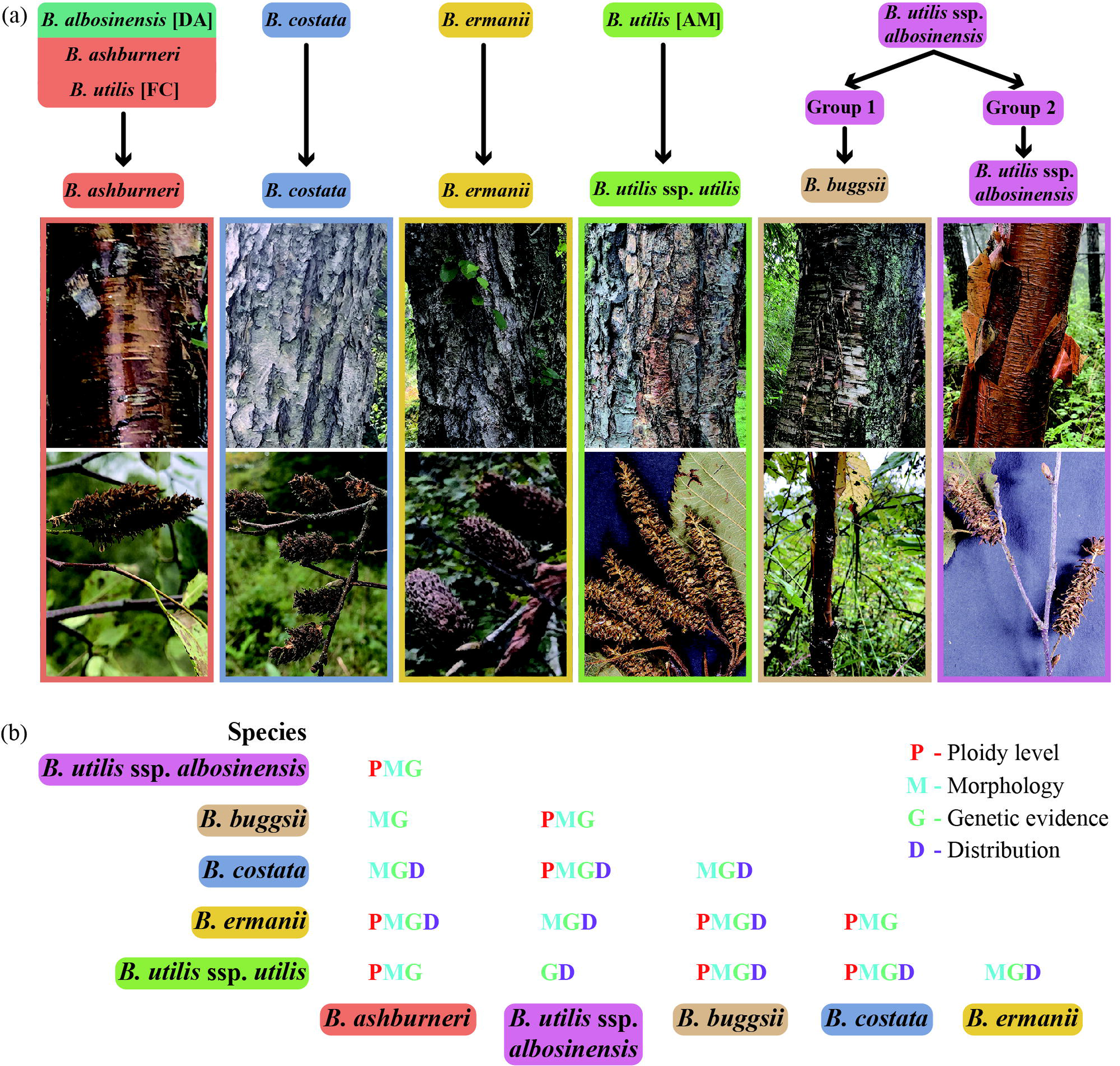
A schematic illustration of species delineation within section *Cosatate* (a) and various sources of information used to distinguish species (b). Photos of each species were placed below its names.

###### Type

CHINA. Chongqing: Chengkou County, elev. ca. 1600-2000 m, 108.7 E, 31.9 N, 6 October 2018 (holotype xx; isotypes xx).

###### Distribution and habitat

*Betula buggsii* occurs in Chongqing, western Hubei and Shaanxi with five localities discovered. *Betula buggsii* grows in mixed forests with bamboos at an altitude of between 1500 and 2100 meters. At some localities, *B. buggsii* grows in parapatry with *B. luminifera* but at a higher altitude. We only founded a small number of *B. buggsii* individuals within each population. Given this situation, we think *B. buggsii* needs conservation.

###### Etymology

*Betula buggsii* is named after Prof. Richard J.A. Buggs, an evolutionary biologist from the Royal Botanical Gardens Kew and Queen Mary University of London, for his devotion to research on hybridisation, phylogenetics and conservation of the genus *Betula*. The Chinese name of *B. buggsii* is “年桦” (nián huá).

## Discussion

### Species delimitation within section *Costatae*

Here we have combined genetic, morphological and distribution data to revise species delimitation within section *Costatae* (genus *Betula*). Our results support six genetic units and thus prefer recognition of six taxa.

### Cluster one — *B. albosinensis* [DA], *B. ashburneri* and *B. utilis* [FC]

Several lines of evidence jointly support the merging of *B. albosinensis* [DA], *B. ashburneri* and *B. utilis* [FC]. First, PCO and STRUCTURE analyses of microsatellite markers indicate an indistinguishable cluster of *B. albosinensis* [DA] and *B. utilis* [FC] (Figs. 3a, 4a). This was further corroborated by admixture analysis of SNPs, showing an indistinguishable cluster of *B. albosinensis* [DA] and *B. utilis* [FC] (Fig. 4b). However, admixture analysis of SNPs including *B. ashburneri* shows the same genetic cluster of *B. ashburneri* and *B. utilis* [AM] (Fig. 4b). By contrast, phylogenomic analysis based on a much larger number of SNPs shows a fully-supported monophyletic clade of *B. albosinensis* [DA], *B. ashburneri* and *B. utilis* [FC] (Fig. 5). The genetic similarity between *B. ashburneri* and *B. utilis* [AM] based on admixture analyses of SNPs suggests *B. ashburneri* being a recent parent of *B. utilis* [AM]. This has been confirmed based on a recent phylogenomic analysis (Wang et al., 2020). In addition, gene flow between the two species may further result in genetic similarity. *Betula ashburneri* is diploid based on chromosome number and genome size analysis (Ashburner & McAllister, 2016; Wang et al., 2016), consistent with the observation that *B. albosinensis* [DA] and *B. utilis* [FC] are also diploids based on microsatellite markers. This is different from descriptions in Flora of China that *B. utilis* [FC] was a tetraploid (Li & Skvortsov, 1999). *Betula ashburneri* was described to occupy a higher altitude than *B. utilis* [AM] (McAllister & Rushforth, 2011), consistent with *B. utilis* [FC] or *B. albosinensis* [DA] occupying a higher altitude than *B. utilis* ssp. *albosinensis* according to our field observations. In addition, *B. ashburneri* was discovered from SE Tibet and reported to distribute in Sichuan and Shaanxi provinces, overlapping with the distribution of *B. utilis* [FC] and *B. albosinensis* [DA]. Based on these, we think *B. utilis* [FC], *B. albosinensis* [DA] and *B. ashburneri* refer to the same species. *Betula ashburneri* was described to have a multi-stemmed shrubby habit and grow up to four meters in height. However, according to our field observations, it can reach 35 meters in height, consistent with descriptions from Flora of China.

### Cluster two — *B. costata*

Both microsatellite and SNPs indicate that *B. costata* is genetically different from other species of section *Costatae* (Figs. 3-4). Despite the fact that *B. costata* and *B. ermanii* co-occur in some populations, the two are morphologically different in fruit, leaf and bark color. In addition, *B. costata* is a diploid and occupies a lower altitude than *B. ermanii,* which is a tetraploid.

### Cluster three — *B. utilis* [AM]

Despite occupying a morphological continuum with *B. utilis* ssp. *albosinensis* group I, molecular results support *B. utilis* [AM] as a genetically distinct unit. *Betula utilis* [AM] are described from the Himalayas, northwestern Yunnan and with an extension into western Sichuan where it coexists with *B. utilis* ssp. *albosinensis* group I.

### Cluster four — *B. utilis* ssp. *albosinensis* group I

*Betula utilis* ssp. *albosinensis* group I forms a morphological continuum with *B. utilis* [AM]. However, molecular analyses indicate that *B. utilis* ssp. *albosinensis* group I forms a distinct cluster with *B. utilis* [AM] (Fig. 4b). We also found that two individuals of *B. utilis* ssp. *albosinensis* group I (XLA01 and XLA32), collected from its northern distribution, show a genetic admixture between *B. utilis* ssp. *albosinensis* and *B. ermanii* (Fig. 4b), indicating their hybrid origin. The two individuals were close to the southern distribution of *B. ermanii,* making hybridisation potentially occur due to long-distance transportation of pollen by wind. In addition, the previously described *B. albosinensis* var. *septentrionalis* and *B. utilis* var. *prattii* are more genetically similar to *B. utilis* ssp. *albosinensis* group I; however, the previously described *B. utilis* ssp. *albosinensis* is genetically similar to *B. utilis* [AM] (Fig. 4b). This indicates some misidentification of these taxa. This was suggested by observations on the very limited number of provenances in cultivation in the UK which led Ashburner and McAllister to describe these taxa as subspecies. Interestingly, the included *B. albosinensis* var. *septentrionalis*, *B. utilis* var. *prattii* and *B. utilis* ssp. *albosinensis* are from Sichuan province where *B. utilis* [FC] and *B. utilis* ssp. *albosinensis* were reported to co-occur. Great morphological variations exist within some populations in Sichuan according to our field observations that even bark color within population shows substantial variation. This made assigning individuals there to either *B. utilis* ssp. *albosinesis* group I or *B. utilis* [AM] impossible based solely on morphological characters.

### Cluster five —*B. ermanii*

*Betula ermanii* and *B. utilis* ssp. *albosinensis* group I are genetically similar based on microsatellite markers but genetically distinct based on SNPs. This is possibly due to very recent gene flow between *B. ermanii* and *B. utilis* ssp. *albosinensis* group I. However, here we think *B. ermanii* should be recognised as a genetic unit on grounds of morphological characters and distribution. Morphologically, *B. ermanii* shows apparent differences in fruit, leaf shape, bark color and the pattern of bark peeling with *B. utilis* ssp. *albosinensis* group I. Geographically, *B. ermanii* distributes around the Changbai Mountains and its north in northeast China where *B. utilis* ssp. *albosinensis* group I is absent there.

### Cluster six — *B. utilis* ssp. *albosinensis* group II (*B. buggsii* as discussed below)

Our genetic analyses revealed a distinct cluster of *B. utilis* ssp. *albosinensis* (group II), which was sufficiently differentiated when compared to other taxa in the genus to be ranked as a new diploid species of section *Costatae*. Based on multiple lines of evidence we describe this new species as *B buggsii*. Molecular analyses of microsatellite markers and SNPs show that *B. buggsii* is genetically distinct from all the other species of section *Costatae*. Phylogenetic analysis based on ITS sequences shows that *B. buggsii* samples cluster together despite low support values (Supplementary Data, Fig. S6). Furthermore, phylogenomic analysis including nearly genus-wide diploid species shows a fully supported monophyletic clade of *B. buggsii,* which was placed within section *Costatae* (Fig. 5). This allows us to confidently establish *B. buggsii* as a new species of section *Costatae*. Interestingly, microsatellite markers revealed two alleles at heterozygous sites for *B. buggsii* whereas three or four alleles for *B. utilis* ssp. *albosinensis,* suggesting a difference in ploidy level. Apart from these, *B. buggsii* shows morphological difference with *B. utilis* ssp. *albosinensis* in bark color and the patterns of bark exfoliation (Fig. 6a). *Betula buggsii*’s bark color is light brown and exfoliates along the elongated lenticels in stripes while *B. utilis* ssp. *albosinensis’s* bark is red and exfoliates in large sheets or flakes (Fig. 6a). The overall morphological similarity between *B. buggsii* and *B. utilis* ssp. *albosinensis* supports the placement of *B. buggsii* within section *Costatae*. Unfortunately, we failed to obtain fruiting catkins, however, we observed seedlings of *B. buggsii* in open habitats, indicating its ability to regenerate and its regeneration depends on habitat disturbance like *B. utilis* ssp. *albosinensis* (Guo et al., 2019).

### A framework for species delimitation within section *Costatae*

A combination of various sources of information (i.e. genetic data, morphological characters, ploidy level and geographic origins) facilitates demarcating species within a morphological or genetic continuum (Fig. 6b). For example, ploidy level is useful in distinguishing a species complex of differing ploidy levels. Recognition of cytotypes would help for conservation purposes as different cytotypes may have different adaptive potentials and are often genetically differentiated. If species reveals a morphological continuum, genetic data and geographic origins would help for distinguishing. This is just the case for *B. utilis* [AM] and *B. utilis* ssp. *albosinensis* group I. Similarly, for species which occupy a genetic continuum, such as *B. utilis* ssp. *albosinensis* I and *B. ermanii,* both morphological data and geographic origin aid in identification..

Finally, for the challenging tetraploids within section *Costatae,* we propose that the most practical taxonomy is to treat populations in north-west Yunnan and the eastern and central Himalaya as *B. utilis* ssp. *utilis;* those from the Qinling Mountains as *B. utilis* ssp. *albosinensis;* and those from northeastern China (e.g. Changbaishan) as *B. ermanii*. For the diploids, it is reasonable to recognise *B. ashburneri, B. buggsii* and *B. costata* based on genetic data, morphological characters and geographic origins. The tetraploids certainly hybridise in cultivation (obser.) and so are likely to hybridise where they co-occur in the wild, generating intermediates with various levels of genetic admixture. Hence, populations collected from region between northwestern Yunnan and the Qinling Mountains may be hybrids between *B. utilis* ssp. *utilis* and *B. utilis* ssp. *albosinensis* and these from between Hebei and northeast China, may be hybrids between *B. utilis* ssp. *albosinensis* and *B. ermanii*. Further research is needed to characterise patterns of genetic admixture between these species within their geographic distributions and to guide future management of genetic diversity.

## Acknowledgements

This work was funded by the National Natural Science Foundation of China (31770230 and 31600295) and Funds of Shandong ‘Double Tops’ Program (SYL2017XTTD13).

---

**Figure S1** Principal coordinate analysis (PCO) of section *Costatae* species at 15 microsatellite markers (a) and principal component analysis (PCoA) of section *Costatae* species at 82,137 SNPs (b).

**Figure S2** The best number of clusters inferred using “Evanno test” method.

**Figure S3** STRUCTURE results of section *Costatae* at K values from 2 to 6 based on 15 microsatellite markers.

**Figure S4** The cross-validation error for each K value from 1 to 10.

**Figure S5** Admixture results at K values from 2 to 10 based on 82,137 SNPs.

**Figure S6** Phylogenetic tree from the maximum likelihood analysis of *B. buggsii* using ITS sequences. Species were classified according to Ashburner and McAllister (2016). Values above branches are bootstrap percentages of >50 %.

## Table legends

**Table S1** Detailed information on populations used in the present study.

**Table S2** Details of microsatellite primers used in the present study.

**Table S3** Detailed information of samples used for ITS and RAD sequencing.

